# Formate, hydrogen, and acetate shape the morphology, physiology and lifestyles of the host-associated methanogen *Methanobrevibacter smithii*

**DOI:** 10.64898/2026.11.14.766705

**Authors:** Isis Del Mazo-Monsalvo, Alexander Neil Poulter, Cesar Ivan Ovando-Ovando, Michel Geovanni Santiago-Martínez

## Abstract

Methane-producing archaea (methanogens) are increasingly linked to gastrointestinal, neuropsychiatric, and neurodevelopmental disorders. These conditions are often associated with altered microbial metabolism that changes the availability of acetate, formate, hydrogen (H_2_), carbon dioxide (CO_2_), and methane (CH_4_) in the gut. However, the physiological importance of formate and acetate in host-associated methanogens remains poorly understood, limiting our ability to interpret microbiome data and infer functions and cross-domain interactions of methanogens in health and disease. Here, we investigated the physiological and metabolic capabilities of *Methanobrevibacter smithii*, the predominant methanogen of the human gut. Formate limitation altered growth, morphology, extracellular pH, gas metabolism, and cellular organization, and induced the formation of cell aggregates. Although acetate was consumed only in small quantities (nmol), its availability significantly influenced growth, gas metabolism, extracellular pH, and formate utilization, indicating functions that extend beyond its role as a biosynthetic precursor. Conversely, nutrient excess promoted a transition to a biofilm lifestyle, suggesting that biofilm formation may function as a carbon sink. We further developed Msm_MEP, an experimentally refined genome-scale metabolic model based on the AGORA2 reconstruction, which captures the metabolic capabilities of M. smithii, quantifies ATP allocation across cellular processes, and predicts the phenotypic consequences of gene deletions. Together, our findings identify formate, hydrogen, and acetate as interacting determinants of *M. smithii* physiology and suggest metabolite-dependent transitions between planktonic growth, cell aggregation, and biofilm formation. Our results establish a framework for interpreting microbiome data and understanding how environmental metabolites shape archaea physiology and cross-domain interactions within the human microbiome.

**Importance:** *Methanobrevibacter smithii* is the predominant methane-producing microorganism (methanogen) in human-associated microbiomes, primarily in the gut; however, its biology remains poorly understood, as it grows slowly and produces little biomass. We show that three common products of microbial metabolism (formate, hydrogen, and acetate) do more than support methane production: they also shape whether *M. smithii* cells divide, aggregate, or form biofilms. Furthermore, methane production is not a reliable indicator of cell growth, which challenges a common assumption in research on methanogens and the human microbiome. By combining laboratory experiments with metabolic models, we discovered that formate and acetate play key roles in shaping cellular functions. Our findings provide a framework for interpreting human microbiome data and predicting how methanogens respond to nutrient fluctuations and interact with bacteria and host cells, thereby advancing our understanding of the roles of methanogens in human health and their relevance to microbiome-based diagnostic and therapeutic strategies.

## Introduction

The human gastrointestinal tract is a densely populated anaerobic ecosystem in which microbial metabolism produces short-chain fatty acids (*e.g.*, acetate, propionate, butyrate), formate, lactate, hydrogen (H_2_), carbon dioxide (CO_2_), methane (CH_4_) and other metabolites as part of nutrient cycling. These metabolites support extensive trophic interactions among microorganisms and can influence intestinal motility, epithelial and immune functions, and communication along the gut-brain axis in the host (Dalile et al., 2019; Pimentel et al., 2006; Samuel, & Gordon, 2006). Alterations in these metabolic networks and microbial community composition (dysbiosis) have been associated with a wide range of disease states across human populations.

Methanogens (anaerobic archaea that produce methane through methanogenesis) occupy a terminal position in microbial metabolic networks by scavenging fermentation products, primarily H_2_ and CO_2_, and converting them into methane (Buan et al., 2018; Lyu et al., 2018; Catlett et al., 2022). Through this metabolic activity, host-associated methanogens can modify the concentrations of H_2_, CO_2,_ and CH_4_ in the gut, reshape the thermodynamic landscape of bacterial fermentation, and ultimately influence microbial community structure and function.

*Methanobrevibacter smithii* is the most prevalent and abundant methanogen in the human gut (Dridi et al., 2009; Chibani et al., 2022; Malat et al., 2024; Moissl-Eichinger et al., 2026), and its genome encodes metabolic traits that may facilitate cross-feeding interactions, microbial community assembly, and host colonization, such as adhesin-like proteins (Samuel et al., 2007; Gupta & Seedorf 2024; Tan et al., 2026). Consistent with these ecological roles, increased methane production and enrichment of *M. smithii* have been associated with gastrointestinal disorders, including constipation, inflammatory bowel disease, and intestinal methanogen overgrowth (IMO) (Hoegenauer et al., 2022; Dirks et al., 2025; Mohammadzadeh et al., 2026). More recently, alterations in archaeal community structure and increased abundance of *M. smithii* have also been linked to colorectal cancer, schizophrenia, and autism spectrum disorder (ASD) (Dirks et al., 2025; Su et al., 2024; Zhu et al., 2025; Fu et al., 2026). These associations have been accompanied by alterations in the metabolism of short-chain fatty acids, amino acids, and formate and other one-carbon compounds, suggesting that cross-domain interactions may influence host health.

Previous culture-based studies have shown that, by consuming bacterial fermentation products, *M. smithii* can alleviate feedback inhibition of bacterial metabolism and influence the flow of energy and nutrients within the gut ecosystem (Samuel et al., 2007; Ruaud et al., 2020; Catlett et al., 2022; Wang et al., 2023). These, co-culture experiments have demonstrated the existence of syntrophic interactions between *M. smithii* and fermentative gut bacteria, involving cross-feeding and the interspecies transfer of H_2_, CO_2_ and other metabolic products. Consequently, H_2_ and CO_2_ are generally considered the principal substrates for methanogenesis of *M. smithii*, whereas acetate serves as a carbon source for biosynthesis (Catlett et al., 2022). Formate is also commonly assumed to substitute for H_2_ and/or CO_2_ during methanogenesis, as reported for several free-living methanogens (Schauer & Ferry, 1980; Holden & Sistu, 2023; Casini et al., 2023; Halim et al., 2024). As a result, fluctuations in the availability of these metabolites are frequently used to interpret human microbiome data. However, direct experimental measurements linking nutrient availability to methanogen physiology remain scarce, limiting our ability to explain the often-observed disconnect between the abundance of archaea and methane emission in humans and other mammals (Kumpitsch et al., 2026).

Here, we investigated how formate, acetate, and methanogenesis-related gases shape the growth, metabolism, morphology, and cellular organization of *Methanobrevibacter smithii*. To mechanistically interpret these phenotypes, we refined the AGORA2 genome-scale metabolic model (GEM) of *M. smithii* (Heinken et al., 2023) using our experimental measurements and published physiological data. The resulting model, Msm_MEP, captures the metabolic capabilities of *M. smithii*, quantifies ATP allocation across cellular processes, and predicts the phenotypic consequences of gene deletions that cannot currently be tested experimentally because genetic tools for this organism are not yet available. By enabling *in silico* interrogation of key genes, reactions, and metabolic pathways, Msm_MEP provides a transferable platform for studying the physiology of methanogens. Because the model is built upon conserved genes and metabolic pathways, it may also provide a foundation for developing and refining genome-scale metabolic models of other *Methanobrevibacter* strains and closely related host-associated methanogens.

Our results identify formate, hydrogen, and acetate as interacting determinants of *M. smithii* physiology and show metabolite-dependent transitions between planktonic growth, cellular aggregation, and biofilm formation. More broadly, our findings suggest that the physiological state of *M. smithii*, rather than its abundance alone, may be a critical determinant of its ecological functions within the human gut. By linking nutrient availability to methanogen physiology, energy allocation, and cellular organization, this work provides a framework for interpreting methanogen-associated signatures in microbiome datasets and for understanding cross-domain interactions among archaea, bacteria, and host cells.

## Results

### Nutrient availability shapes the cell morphology and lifestyles of *M. smithii*

To investigate how nutrient availability influences its physiological and metabolic capabilities, we developed a cultivation platform for *Methanobrevibacter smithii* PS (DSM 861; ATCC 35061), the type strain (Miller et al, 1982) and an emerging model for host-associated methanogens.

We formulated MEP01, a defined oxygen-free liquid medium that enables systematic evaluation of physiological responses to methanogenic substrates and nutrients. MEP01 contains an 80% H_2_/20% CO_2_ headspace and is supplemented with 22 mM acetate and 30 mM formate in the liquid phase (***See* Methods**). Cultures were incubated at 37°C with agitation (110 rpm) in the dark. The composition of MEP01 was based on previously described media for *M. smithii* but excludes yeast extract, tryptone, and other undefined components, permitting precise manipulation of nutrient availability (Miller et al., 1982; Pende et al., 2021; Catlett et al., 2022; Pilliol et al., 2023; Weinberger et al., 2025a). Under the conditions used in this study, bicarbonate was an essential component and could neither be omitted nor replaced by other buffering agents (Santiago-Martínez & Ferry, 2023).

Cultures of *M. smithii* exhibited a pearly-white appearance during active growth and methanogenesis (**Fig. 1A-B**), gradually transitioning to a yellowish coloration upon entry into stationary phase. Consistent with previous reports, cultures appeared uniformly turbid due to suspended cells (Miller et al., 1982; Pende et al., 2021; Catlett et al., 2022; Weinberger et al., 2025a; Smith et al., 2026). However, dispersed microcolonies were also visible as fine particulate structures, indicating that cellular organization was not entirely homogeneous. As nutrients became limiting during late stationary phase, these cultures transitioned to an aggregative phenotype characterized by small cellular clusters (cell aggregates) and a macroscopically heterogeneous suspension (**Fig. 1D**).

**Figure 1.**
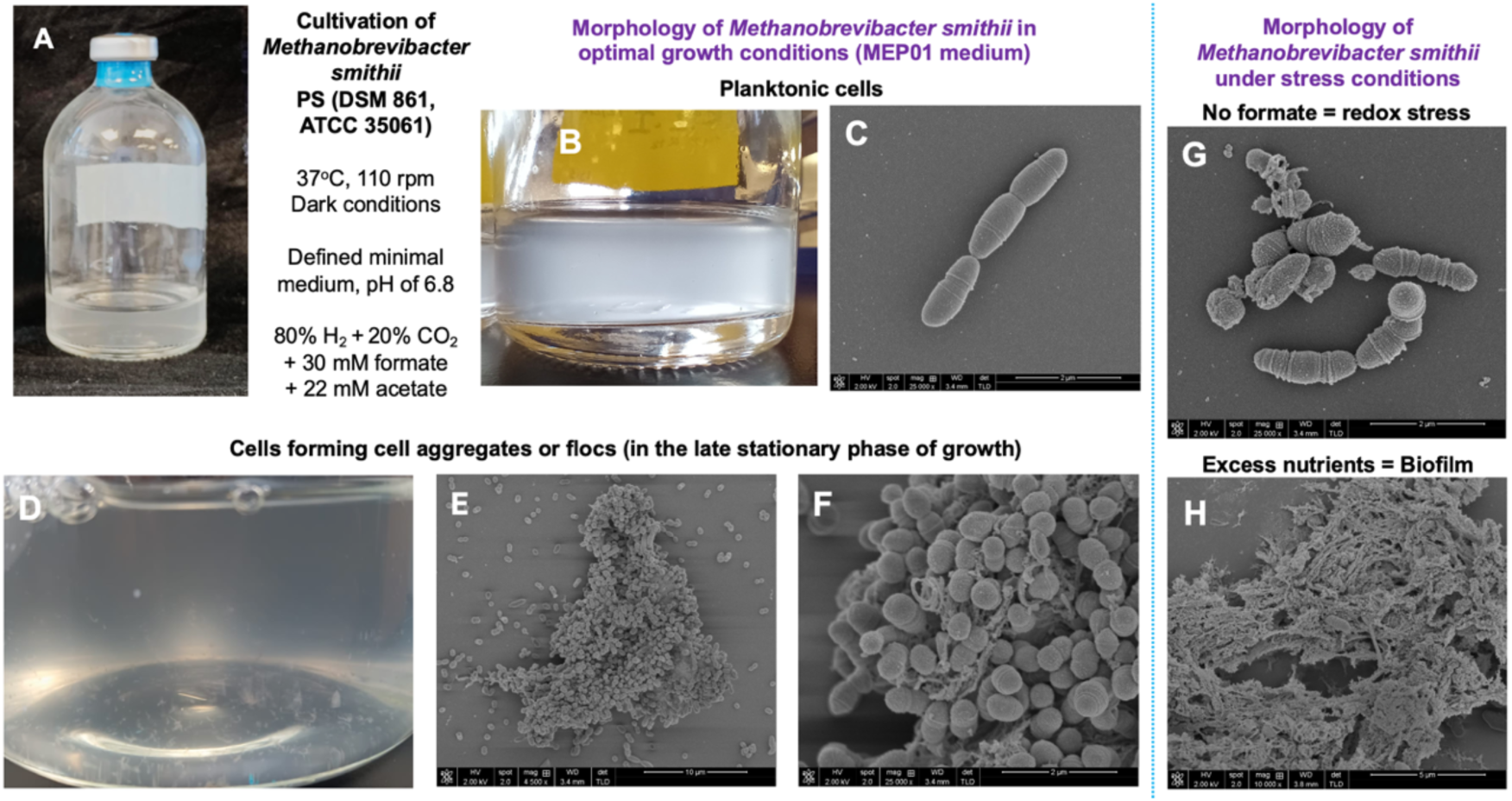
Nutrient availability shapes the morphology and lifestyles of *Methanobrevibacter smithii*. (A) Cells were cultured anaerobically in 118-mL serum bottles containing 20 mL of oxygen-free liquid medium. A constant liquid-to-headspace ratio was maintained to support reproducible growth. (B) During the active growth phase, cells exhibit a planktonic lifestyle and remained suspended in the medium. (C) Cells displayed an ovococcoid, nonmotile morphology, occurring predominantly as pairs or short chains in scanning electron microscopy (SEM) images. (D) During the late stationary phase of growth, cells formed macroscopic aggregates, and (E, F) appeared as compact multicellular clusters in SEM images. (G) Cells exposed to nutrient stress (no formate) exhibited greater variation in shape and size, (H) whereas nutrient excess promoted biofilm formation (H).

During the active growth phase, the cells only exhibit a planktonic lifestyle. Cells exhibited an ovococcoid, nonmotile morphology, measured approximately 1 µm in length, and occurred predominantly as pairs or short chains in scanning electron microscopy (SEM) images (**Fig. 1C**). In contrast, in the late stationary phase, cells formed macroscopic flocs that appeared by SEM as compact multicellular aggregates (**Fig. 1D-F**). The absence of formate increased morphological heterogeneity (**Fig. 1G**), whereas nutrient excess (*e.g.*, substrates, vitamins, metals) promoted extracellular matrix production and the development of a biofilm-like growth lifestyle (**Fig.1H**).

The cell aggregates (**Fig. 1D-F**) were resistant to gentle agitation (100 rpm) but readily dispersed under vigorous shaking. Treatment with Triton X-100, DNase I, or proteinase K did not induce aggregate disassembly. However, transfer of late-stationary-phase aggregates to fresh MEP01 medium partially restored planktonic growth, indicating that a subset of cells remained viable and that the aggregative phenotype could be reversible and responsive to nutrient availability.

### *Methanobrevibacter smithii* exhibits slow growth, low biomass yield, and the transition to an aggregative lifestyle

To evaluate the effect of inoculum size on growth, cultures were initiated at Optical Density at 600 nm (OD_600_) values ranging from 0.020 to 0.070. This range was selected based on preliminary cultivation studies and previous experience with methanogenic archaea. An intermediate inoculum (OD_600_ = 0.035–0.040; approximately 6 mL culture) was chosen for subsequent experiments because it provided the greatest dynamic range for detecting treatment effects while maintaining consistent and reproducible growth kinetics across more than 30 independent experiments (**Fig. 2B**).

**Figure 2.**
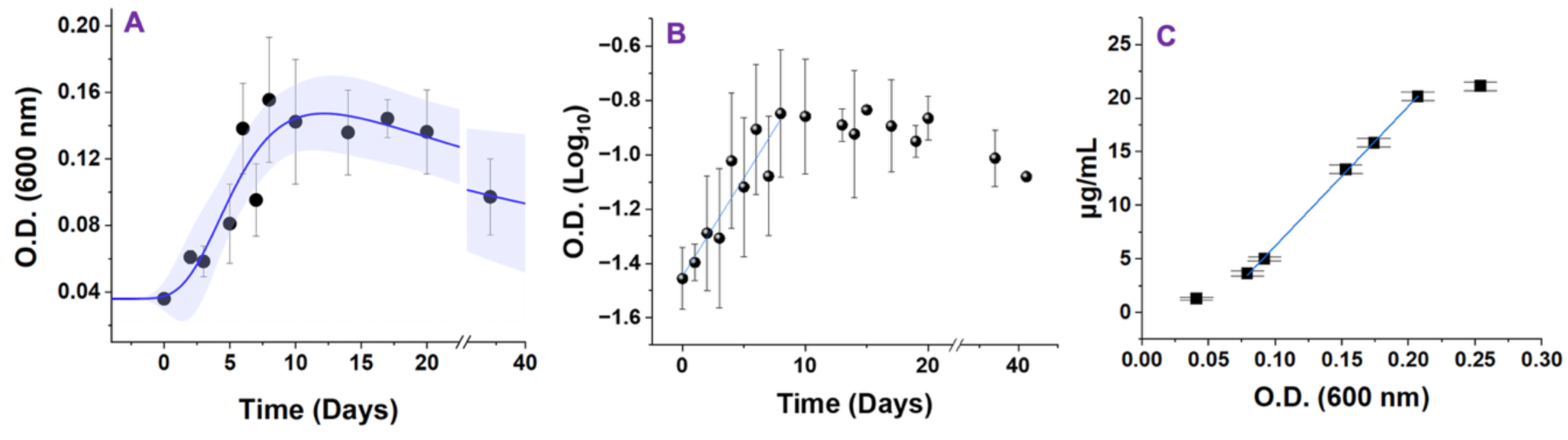
Methanobrevibacter smithii grows slowly and produces low biomass in MEP01 media. (A) Growth monitored by optical density at 600 nm (OD_600_). (B) Semi-logarithmic growth curve showing exponential growth during the first 10 days, followed by the onset of cell aggregation, more evident after day 20. (C) Relationship between OD_600_ and total abundance of protein, used as an independent measure of biomass accumulation and cell growth. The values represent the mean ± standard deviation of 30 independent experiments in A and B, and of 4 independent experiments in C.

In all subsequent experiments, a constant liquid-to-headspace ratio was maintained (20 mL medium in 118-mL serum bottles; **Fig. 1A**) to ensure reproducible growth and facilitate comparisons across conditions. A detailed description of the inoculation procedure is provided in the Supplementary Materials.

To determine which model best described the growth dynamics of *M. smithii* (**Fig. 2A**), we compared four equation-based growth models (Zwietering et al., 1990): the Gompertz model, a four-parameter logistic model, a sigmoidal model previously applied to methanogens, and our modified Gompertz model incorporating an exponential decay term to account for cell aggregation. Additional models, including biphasic functions, were evaluated but either produced biologically unrealistic parameter estimates or yielded large uncertainties. Consequently, subsequent analyses were restricted to these four models summarized in **Table 1**. The corresponding equations are described in the Methods section.

**Table 1.** Growth model parameters describing the growth dynamics of the host-associated methanogen *Methanobrevibacter smithii*.

| **Model** | **R<sup>2</sup>** | **Reduced Chi<sup>2</sup>** | **Key parameters** |
|---|---|---|---|
| Gompertz | 0.8079 | 2.94×10<sup>-4</sup> | a = 0.133, k = 0.319 |
| Logistic | 0.8110 | 2.89×10<sup>-4</sup> | A<sup>2</sup> = 0.132, p = 3.34 |
| Sigmoidal | 0.8213 | — | K = 0.131, r = 0.449 |
| Our modified Gompertz | 0.827 | 2.24×10<sup>-4</sup> | A = 0.143, δ = 0268 |

Growth of *M. smithii* under standard conditions (MEP01 medium; initial OD_600_ = 0.035) was best described by a modified Gompertz model (R^2^ = 0.827; **Fig. 2A**). Cultures reached a maximum OD_600_ of 0.167 ± 0.054, corresponding to approximately 2.2 cell duplications. The estimated growth rate constant was 0.334 ± 0.135 day^-1^, yielding a maximum specific growth rate (μ) of 0.021 day^-1^, whereas the lag phase lasted 1.62 days. The model also identified a post-stationary decline phase (δ = 0.0268 ± 0.0152 day^-1^), consistent with a gradual loss of apparent biomass due to cell aggregation and/or cell death. Together, these parameters establish a baseline for subsequent physiological comparisons.

Semilogarithmic growth curves indicated exponential growth during the first 10 days of cultivation, followed by the onset of cell aggregation, which became increasingly pronounced after day 20 (**Fig. 2B**). Total cellular protein was quantified as an independent measure of biomass accumulation and displayed a similar overall growth pattern (**Fig. 2C**).

The sampling frequency influenced measurements of growth kinetics and gas metabolism, depending on the number of samples taken along the growth curve. This is likely due to gas loss and changes in the ratio between the liquid phase and the headspace. Nevertheless, the general trends remained consistent, regardless of the sampling regimen (See supplementary material).

### Formate and hydrogen limitation remodel growth, metabolism, and cellular organization in *M. smithii*

#### Standard condition

Under standard culture conditions (MEP01 medium containing an 80% H_2_/20% CO_2_ headspace, 30 mM formate, and 22 mM acetate; initial pH 6.8), methane production followed a sigmoidal trajectory that was well described by the Gompertz model (R^2^ > 0.94; **Fig. 3B**). Mean methane accumulation reached 938.8 ± 162.6 μmol CH_4_, with a maximum production rate of 119.3 ± 3.8 μmol day^-1^, indicating robust and reproducible methanogenesis across six independent experiments.

**Figure 3.**
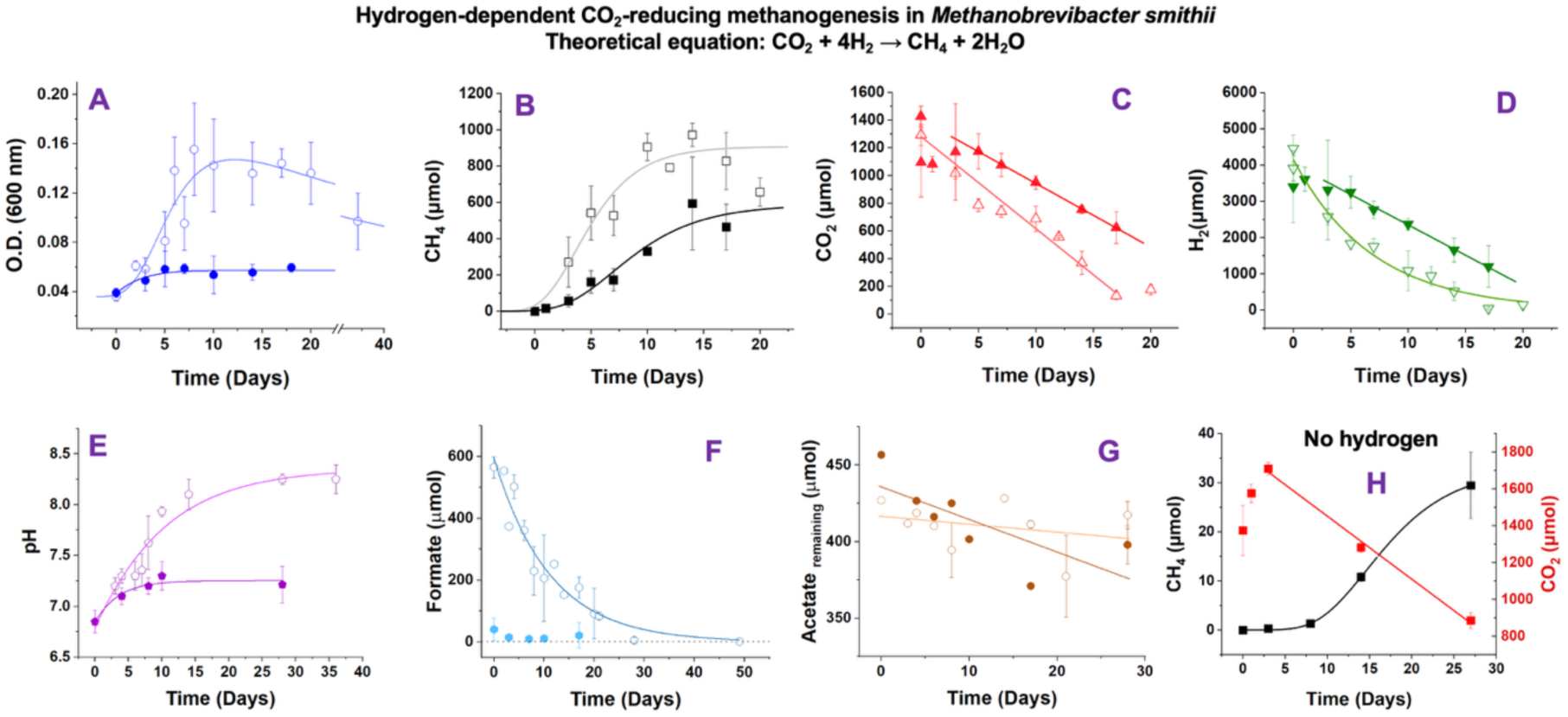
*Methanobrevibacter smithii* performs H_2_-dependent, CO_2_-reducing methanogenesis, and formate availability influences its growth and metabolism. Cultures without formate are represented by closed symbols, whereas cultures containing 30 mM formate are represented by open symbols (complete MEP01 medium). The absence of exogenous formate in the liquid media decreased cell growth (A), methane production (B), carbon dioxide consumption (C), and hydrogen consumption (D) and altered extracellular pH (E). The cells consumed all the exogenous formate (F). A small fraction of the exogenous acetate was consumed, and its consumption rate was also affected by formate omission (G). In the absence of hydrogen (H), the cells produced significantly less methane, indicating that formate did not completely replace hydrogen under the conditions evaluated. Values represent the mean ± standard deviation of 6 independent experiments, except for the standard condition shown in A (n= 30).

Consumption of H_2_ and CO_2_ was consistent with active hydrogen-dependent CO_2_-reducing methanogenesis and was well described by first-order decay models (**Fig. 3C-D**). The initial headspace contained 4.11 ± 0.17 mmol H_2_ and 1.35 ± 0.06 mmol CO_2_, corresponding to an H_2_:CO_2_ ratio of approximately 3:1. During active growth, CO_2_ was removed from the headspace at a nearly constant rate until a stable residual pool remained, whereas H_2_ consumption followed an exponential decay trajectory and approached complete depletion by the end of the experiment.

CO_2_ consumption (**Fig. 3C**) followed a near-linear trajectory during the phase of active growth, although the data were also well described by a first-order decay model (k = 0.092 ± 0.008 day^-1^; t½ = 7.6 ± 0.7 days). During this period, CO₂ declined at a rate of 66.5 ± 5.7 μmol day^-1^ (R^2^ = 0.95) until a stable residual pool of approximately 132 μmol remained in the headspace.

H_2_ consumption (**Fig. 3D**) followed a typical exponential decay pattern (R^2^ = 0.98). Nearly the entire initial H_2_ pool (4109 ± 166 μmol) was consumed, with a rate constant of 0.142 ± 0.012 day^-1^. Together, these results are consistent with active hydrogen-dependent CO_2_-reducing methanogenesis, while the persistence of a residual CO_2_ pool suggests that CO_2_ availability did not become limiting under standard growth conditions.

#### No-formate condition

The absence of formate altered growth (**Fig. 2A**), cellular morphology (**Fig. 1G**), extracellular pH (**Fig. 3E**), gas metabolism (**Fig. 3B-D**), and cellular organization, and promoted cell aggregation. Formate-free cultures reached only 35% of the biomass observed under standard conditions (OD_600max_ = 0.057 ± 0.002 versus 0.164 ± 0.054), indicating that formate is not essential for initiating growth but is required for sustained biomass accumulation. Consistent with this interpretation, growth was best described by the standard Gompertz model (R^2^= 0.864), reflecting an initial growth phase followed by rapid growth arrest.

Methane production was substantially reduced in the absence of formate, with maximum CH_4_ accumulation reaching only 587 ± 124 μmol, a 38% decrease relative to the standard condition (**Fig. 3B**). CO_2_ consumption also exhibited altered kinetics (**Fig. 3C**), shifting from first-order decay under standard conditions to an approximately linear trend during the metabolically active phase. This transition is consistent with a change from substrate-dependent consumption to a rate limited by cellular metabolic capacity. H_2_ consumption remained well described by first-order kinetics but proceeded 46% more slowly in formate-free cultures than under standard conditions (**Fig. 3D**). Despite the slower kinetics, total H_2_ consumption differed slightly between conditions, indicating that formate primarily affected the rate, rather than the extent, of hydrogen utilization.

Formate deprivation also altered extracellular pH dynamics. Whereas the standard condition reached a final pH of 8.34 ± 0.12, formate-free cultures stabilized at a significantly lower pH of 7.25 ± 0.05 (*p* < 0.0001; **Fig. 3E**), indicating substantial changes in carbon and proton fluxes. Acetate concentrations exhibited a tendency to decline more rapidly in formate-free cultures (**Fig. 3G**), although variability prevented definitive conclusions. Given that only a few nanomoles of acetate would be sufficient to account for the observed biomass yield, any net acetate consumption was expected to be small and therefore difficult to quantify, even using a highly sensitive enzymatic assay. These observations suggest that *M. smithii* is more likely to respond to changes in acetate availability than to substantially alter acetate pools within the gut ecosystem.

#### No-hydrogen condition

Replacement of H_2_ with argon drastically reduced methanogenesis and triggered a rapid decline in culture density (**Fig. 3H**). Optical density decreased progressively from 0.040 ± 0.001 to 0.026 ± 0.004 over 27 days, corresponding to a 35% reduction in biomass. This sustained decline indicates that, in the absence of H_2_ as an electron donor, *M. smithii* is unable to maintain a stable metabolic state and instead enters a decay phase.

Biomass loss was accompanied by a pronounced morphological transition. Cells formed macroscopic flocs and progressively sedimented, replacing the finely dispersed phenotype characteristic of actively growing cultures (**Fig. 3H**). These flocs rendered cultures macroscopically heterogeneous, with visible aggregates persisting even after gentle agitation. The formation of large sedimented aggregates under conditions of total H_2_ depletion contrasts with the smaller aggregates observed under nutrient limitation, suggesting that different levels of energy stress result in distinct organizational states in *M. smithii.* Together, these observations indicate that hydrogen availability not only sustains methanogenesis but also influences cellular organization and population structure.

#### Tungsten (W) supplementation

Given the role of tungsten-dependent enzymes (*e.g.*, formylmethanofuran dehydrogenase, formate dehydrogenase, and other oxidoreductases) in hydrogen-dependent CO_2_-reducing methanogenesis (Halim et al., 2024), we also examined the effects of tungsten availability on the growth and physiology of *M. smithii* (**Fig. 4**). Trace tungsten supplementation (0.05 µM W) increased both the rate and total yield of methane production without affecting final biomass accumulation. In contrast, higher tungsten concentrations (0.5 and 5 µM) did not significantly alter methane production or biomass yield but reduced cell aggregation (**Fig. 4C**). Rates of CO_2_ and H_2_ consumption were largely unaffected by tungsten supplementation, although 5 µM W reduced total H_2_ consumption. Together, these results suggest that tungsten is not a major growth-limiting factor under standard conditions, although trace supplementation can enhance methanogenesis and influence cellular organization.

**Figure 4.**
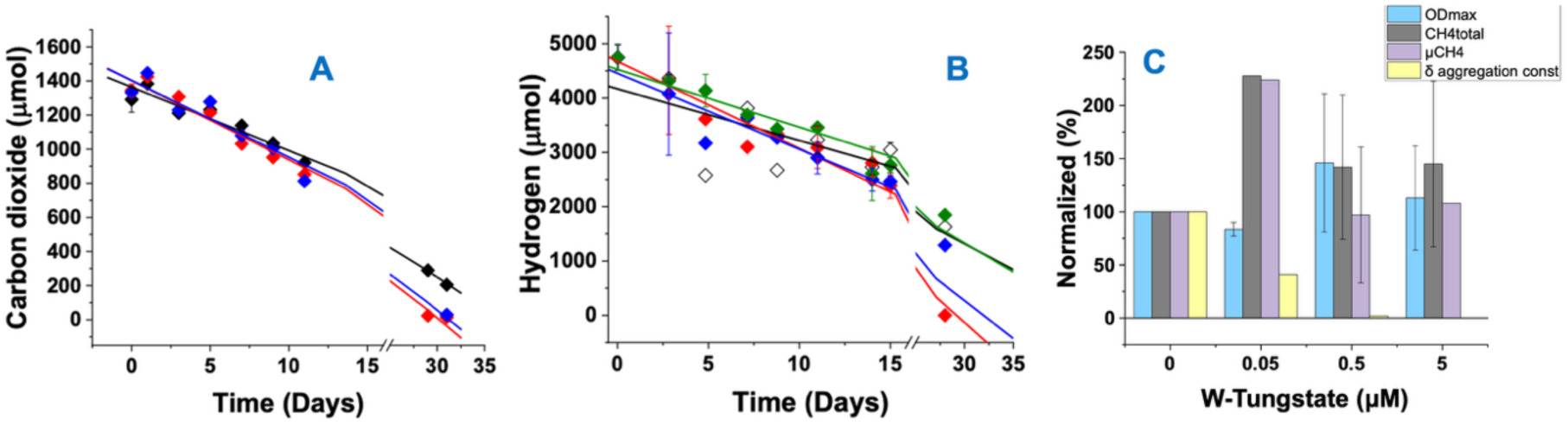
Effect of tungsten (W) supplementation on the growth and metabolism of *Methanobrevibacter smithii*. Tungsten supplementation did not affect the rates of carbon dioxide (A) or hydrogen (B) consumption, although minor differences in final gas consumption were observed after day 20. Black/white diamond = control experiments in which data obtained under standard conditions (MEP01 medium) are represented; red diamond= 0 µM W; blue diamond = 0.05 µM W; green diamond= 0.5 µM W. (C) Effects of tungsten concentration on maximum optical density at 600 nm (OD_600_), total methane production, methane production rate, and the cell aggregation constant. Values represent the mean ± standard deviation of 4 independent experiments.

#### Iron (Fe) supplementation

Because iron is a key component of hydrogenases, electron-bifurcating HdrABC, oxidoreductases, ferredoxins, and other Fe-S proteins involved in methanogenesis and biosynthetic pathways (Wagner et al., 2017; Halim et al., 2024; Bonitatibus et al., 2025), we evaluated the effects of iron supplementation on the growth and physiology of *M. smithii* (**Fig. 4**). Addition of 50 µM Fe^2+^ substantially increased both growth and methanogenesis relative to standard conditions. Maximum biomass increased 2.3-fold (OD_600_ = 0.330 ± 0.036 versus 0.146 ± 0.001; *p* < 0.001), while total CH_4_ production increased 2.2-fold (2462 ± 254 versus 1122 ± 289 µmol). Rates of H_2_ and CO_2_ consumption were 4.4-fold and 5.2-fold higher, respectively, in iron-supplemented cultures.

Despite these effects on culture performance, biomass-normalized methanogenic activity remained nearly identical under both conditions (≈7,500 µmol CH_4_ per OD unit), and formate consumption kinetics were similar (t_1_ = 10.02 ± 3.36 versus 10.54 ± 1.01 days). These results indicate that the primary effect of iron was to promote biomass accumulation rather than to enhance the intrinsic metabolic activity of individual cells.

Together, these findings suggest that iron availability was suboptimal for maximal biomass production under standard conditions, but sufficient to support reproducible growth and preserve the characteristic physiological responses of *M. smithii*. Consequently, the standard MEP01 medium remained appropriate for evaluating metabolite-dependent effects while maintaining sensitivity to detect changes in cell growth, metabolism, and cellular organization.

#### Acetate availability exerts disproportionate effects on growth and metabolism despite limited consumption

Acetate limitation (2 mM instead of 22 mM) substantially impaired growth, gas metabolism, extracellular pH, and formate utilization (**Fig. 5**). Maximum biomass remained low (OD_600_= 0.065-0.069), reaching only ∼40% of that observed under standard conditions despite extended incubation periods. Consistent with this reduction in growth, H_2_ and CO_2_ consumption rates decreased by 20-60%, resulting in an approximately 50% reduction in methane production.

**Figure 5.**
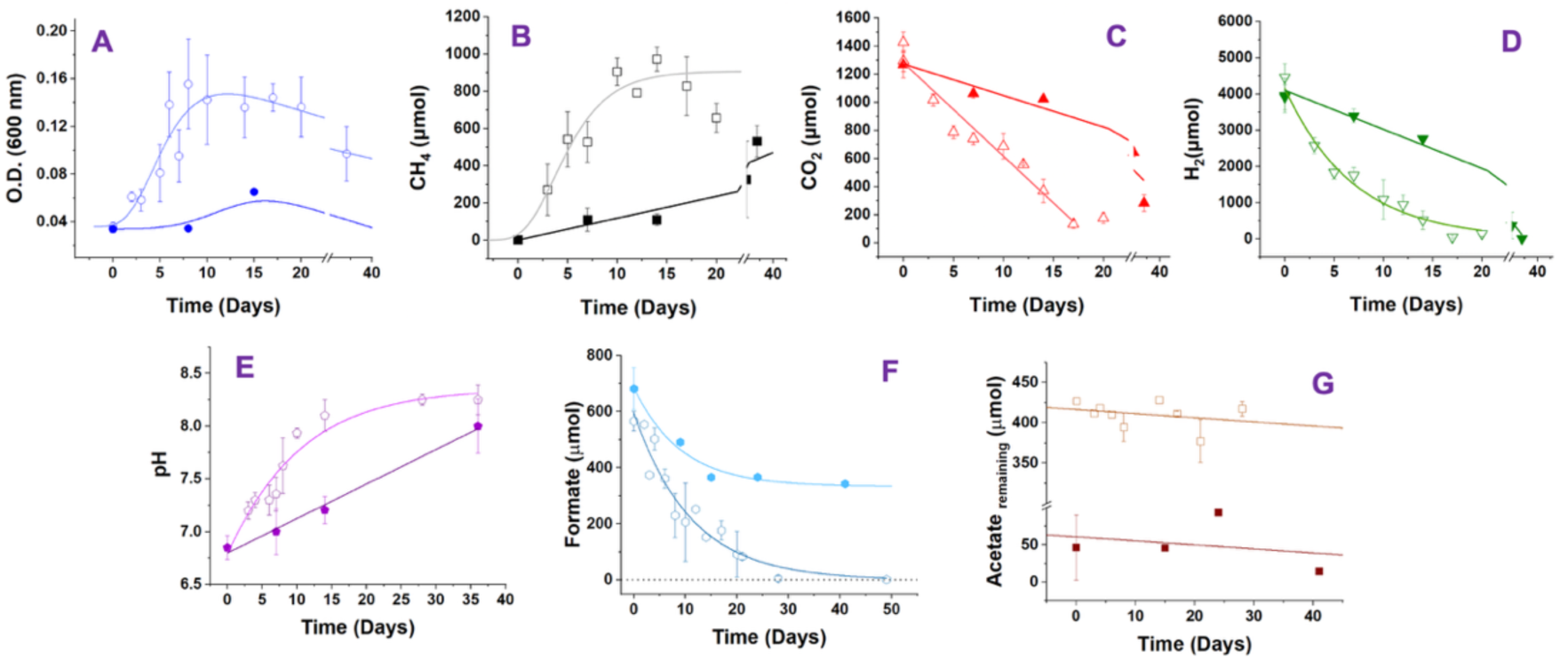
Acetate availability influences the growth and metabolism of *Methanobrevibacter smithii*. Cultures containing 2 mM acetate are represented by closed symbols, whereas cultures containing 22 mM acetate are represented by open symbols. The lower acetate concentration decreased cell growth (A), methane production (B), carbon dioxide consumption (C), and hydrogen consumption (D). Acetate availability also affected the rate of change in extracellular pH, although cultures reached similar final pH values (E). Exogenous formate was only partially consumed (F), whereas no clear net consumption of exogenous acetate (G) was detected under low-acetate conditions. Values represent the mean ± standard deviation of 3 independent experiments, except for the control condition shown in A (n= 30).

Under standard conditions, formate concentrations declined according to a first-order exponential decay model and were completely consumed with a time constant of 11.1 ± 1.2 days (**Fig. 5F**). In contrast, under acetate-limited conditions, formate consumption remained incomplete throughout the experiment, indicating that acetate availability indirectly influences formate utilization and methanogenesis.

Despite an 11-fold difference in initial acetate concentration, the apparent rates of acetate consumption did not differ significantly between standard and low-acetate conditions (−0.513 ± 0.619 versus −0.539 ± 1.286 μmol day^-1^; *p* > 0.05; **Fig. 5G**). Both conditions exhibited negative slopes consistent with acetate uptake; however, the low rates and high variability prevented definitive quantification of acetate consumption. Importantly, only small amounts of acetate (nmoles) would be required to support the observed biomass yields, suggesting that net acetate utilization approached the limits of analytical detection. Nevertheless, acetate availability strongly influenced growth, methane production, extracellular pH, and formate utilization, indicating that its physiological importance extends beyond its role as a biosynthetic precursor.

#### Msm_MEP: A refined genome-scale metabolic model explains the observed phenotypes and quantifies ATP allocation

To investigate the metabolic basis of the observed phenotypes, we developed a refined genome-scale metabolic model (GEM) that integrates our experimental measurements with published physiological data.

We refined the AGORA2 GEM reconstruction of *M. smithii* PS (DSM 861, ATCC 35061) (Heinken et al., 2023). Our resulting GEM model, designated **Msm_MEP**, contains 733 reactions, 752 metabolites, and 508 genes. Compared with the original AGORA2 reconstruction, Msm_MEP incorporates extensive manual curation, including the removal of 52 reactions and the addition of 95 reactions selected to better represent archaeal metabolism, methanogenesis, cofactor biosynthesis, and energy conservation pathways in *M. smithii* (See Table S3 and Supplementary Material).

Removals from the published AGORA2 GEM were mainly bacterial reactions with no counterpart in *M. smithii* including lipoteichoic acid synthesis, ubiquinone and menaquinone biosynthesis, and quinone-dependent respiration as members of the Methanobacteriales do not produce quinones (Becker et al., 2018). For example, the base model included methanophenazine hydrogenase, a membrane-bound hydrogenase that uses methanophenazine as an electron carrier. Because methanophenazine is characteristic of cytochrome-containing methanogens such as *Methanosarcina mazei* and *Methanosarcina barkeri* (Abken et al., 1998; Welte & Deppenmeier 2014; Mand et al., 2018), this reaction is not expected to occur in *M. smithii*.

Additions to the Msm_MEP model primarily consisted of archaeal pathways that were absent or incomplete in the base model, including: (i) biosynthetic pathways for methanogenesis cofactors, such as coenzyme M (HS-CoM), coenzyme B (HS-CoB), cofactor F_420_, and cofactor F_430_; (ii) energy conservation pathways, including the energy-conserving hydrogenases Eha and Ehb and the electron-bifurcating heterodisulfide reductase complex Mvh-HdrABC; and (iii) archaeal-specific reactions replacing bacterial counterparts (Wagner et al., 2017; Grinter & Greening, 2021; Mekki et al., 2026).

Because methanogenesis is the primary source of ATP in *M. smithii*, we used the Msm_MEP model to quantify how methanogenesis-derived ATP is allocated to biomass synthesis and other cellular processes (**Table 2**), excluding Growth-Associated Maintenance (GAM), which comprises ATP expenditures associated with macromolecule polymerization (*e.g.*, proteins, RNA, DNA, and polysaccharides) and the assembly of cellular structures. The Msm_MEP model predicts that acetate activation accounts for the largest fraction of ATP expenditure associated with biomass synthesis. The second largest ATP demand is associated with the carboxylation of pyruvate and gluconeogenesis.

**Table 2.** Msm_MEP model-predicted ATP cost (excluding GAM) of producing 1 g dry cell weight (gDW) in key metabolic processes.

| **Metabolic process** | **mmol ATP per gDW** | **% ATP (excluding GAM)** |
|---|---|---|
| Acetate activation (AMP-forming Acs) | 28.48 | 40.7 |
| Gluconeogenesis / pyruvate carboxylation | 16.46 | 23.5 |
| Nucleotide synthesis | 6.98 | 10 |
| Na<sup>+</sup> used by Eha for ferredoxin reduction<br>(ATP is not produced) | 4.64 | 6.6 |
| Nitrogen assimilation | 4.59 | 6.6 |
| Other biosynthesis | 3.43 | 4.9 |
| Na<sup>+</sup>/H<sup>+</sup> antiport, proton export<br>(ATP is not produced) | 3.08 | 4.4 |
| Amino-acid pathway kinases | 2.24 | 3.2 |
| Total ATP costs | 69.9 | 100 |

Furthermore, by constraining the Msm_MEP model with the experimentally measured growth and CH_4_ production rates, we estimated the ATP remaining after accounting for the energetic costs of biomass synthesis under each culture condition (**Table 3**). This residual ATP represents non-growth-associated maintenance (NGAM), a term used to describe ATP costs that are not allocated to biomass formation, but may instead contribute to cellular maintenance, repair processes, and other physiological functions (Heinken et al., 2023).

**Table 3.** Msm_MEP model-predicted ATP available for non-growth-associated (NGAM) processes under *in vitro* conditions. The growth and CH_4_ production rates were set to values obtained from fits to our experimental data.

| **Condition** | **Time point** | **OD at 600 nm** | **CH<sub>4</sub> (mmol/gDW per h)** | **ATP from CH<sub>4</sub> (mmol ATP/gDW per h)** | **ATP to growth (mmol ATP/gDW per h)** | **NGAM (mmol ATP/gDW per h)** |
|---|---|---|---|---|---|---|
| Standard | Day 3.90 | 0.079 | 29.53 | 12.55 | 1.78 | 10.77 |
| Standard | Day 4.61 | 0.091 | 22.92 | 9.74 | 1.41 | 8.33 |
| Standard | Day 10.00 | 0.145 | 3.53 | 1.5 | 0.13 | 1.37 |
| No-formate | Day 6.94 | 0.056 | 23.99 | 10.2 | 0.09 | 10.1 |
| Low-acetate | Max OD,<br>linear CH<sub>4</sub> | 0.067 | 2.33 | 0.99 | 0.04 | 0.95 |
| No H<sub>2</sub><br>(argon) | Starting OD | 0.04 | 3.12 | 1.33 | 0 | 1.33 |

When modeling the phenotype of an *Msm_0330* deletion mutant, which lacks the AMP-forming acetyl-CoA synthetase (Acs) (**Fig. 6**), the Msm_MEP model predicted a substantial redistribution of ATP allocation, confirming that acetate activation by Acs represents the single largest ATP costs associated with biomass synthesis. These simulations support the conclusion that acetate assimilation imposes a disproportionate energetic burden relative to the amount of acetate consumed. Together with the experimentally observed low rates of acetate consumption, these findings suggest that *M. smithii* minimizes ATP costs on biosynthesis while maintaining methanogenesis, a strategy that may be advantageous in the nutrient-variable environment of the human gut.

**Figure 6.**
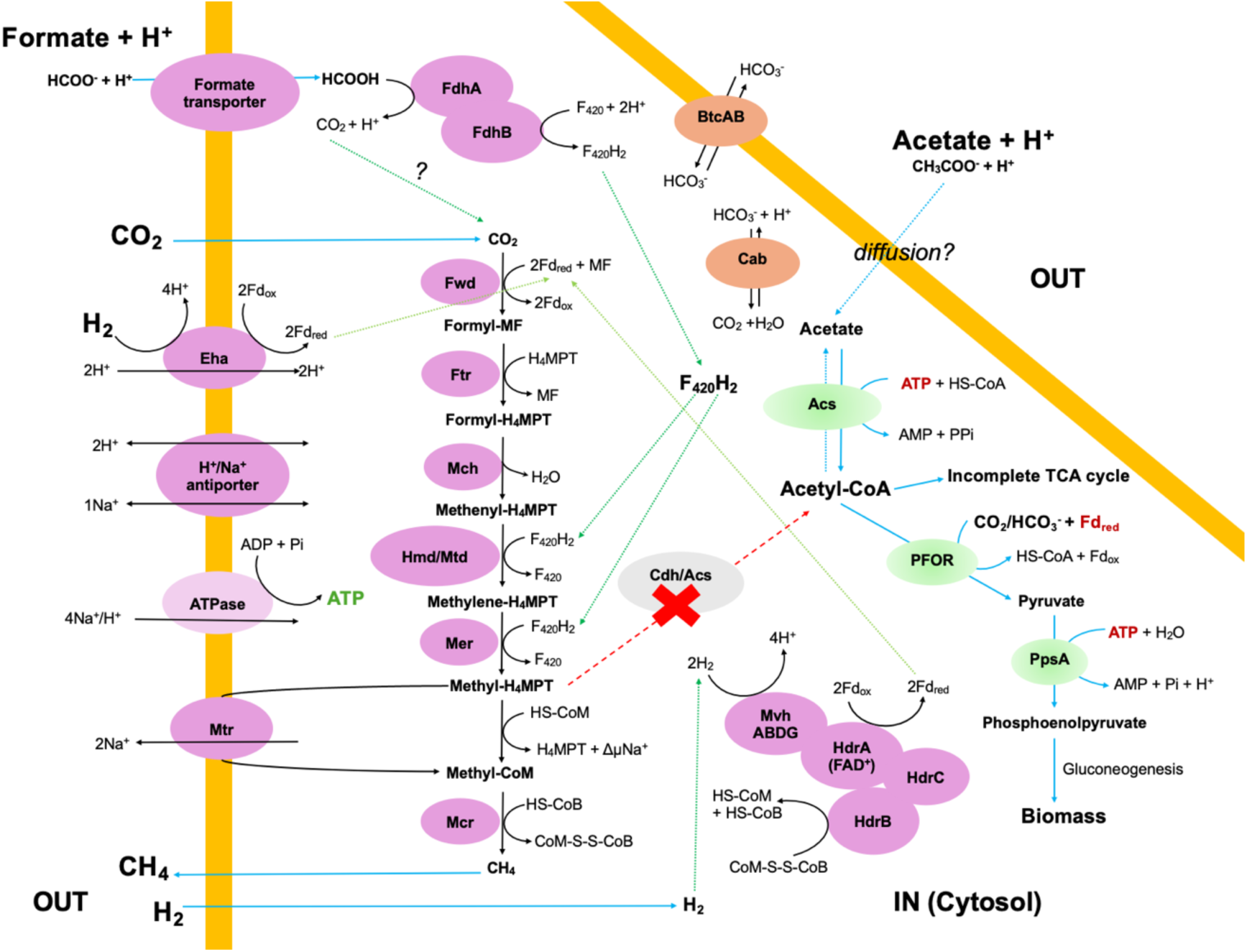
A revised model of hydrogen-dependent CO_2_-reducing methanogenesis in *Methanobrevibacter smithii* that integrates the metabolism of formate and acetate and links methanogenesis to carbon assimilation, energy conservation, and cellular physiology. Methanogenesis is an anaerobic electron transport system in which CoM-S-S-CoB (heterodisulfide) functions as the terminal electron acceptor. Electron transfer, the production of a Na^+^/H^+^ gradient for ATP synthesis, and the interconversion of redox coenzymes involve many steps, and methanogenesis pathways can vary depending on the substrate and electron donors. **Abbreviations:** Acs, acetyl-CoA synthase; ATPase, archaeal A_1_A_0_ ATP synthase; BtcAB, bicarbonate transporter; Cab, carbonic anhydrase; Cdh/ACS, Carbon monoxide dehydrogenase/Acetyl-CoA synthase complex; CH_3_, methyl; Eha, energy-converting hydrogenase; F_420_, cofactor F_420_; Fd, ferredoxin (ox, oxidized; red, reduced); FdhAB, formate dehydrogenase subunits; Ftr, formyl-methanofuran: tetrahydromethanopterin (H_4_MPT) formyl-transferase; Fwd, tungsten formyl-methanofuran dehydrogenase; H_4_MPT, tetrahydromethanopterin; Hmd, H_2_-forming methylene-H_4_MPT dehydrogenase; HS-CoB, coenzyme B; HS-CoM, coenzyme M; Mch, methenyl-H_4_MPT cyclohydrolase; Mcr, methyl-CoM reductase; Mer, methylene-H_4_MPT reductase; Mtd, F_420_-dependent methylene-H_4_MPT dehydrogenase; Mtr, methyl-H_4_MPT:CoM methyltransferase; Mvh-HdrABC, methylviologen-reducing hydrogenase/heterodisulfide reductase complex (an electron bifurcating complex); PFOR, pyruvate:ferredoxin oxidoreductase; PspA, phosphoenolpyruvate synthase.

Overall, our experimental and modeling results show that nutrient availability influences growth, methanogenesis, gas metabolism, formate and acetate utilization, energy allocation, and cellular lifestyle in *M. smithii*. The Msm_MEP model provides a complementary framework for explaining these physiological responses by identifying the metabolic constraints that govern ATP production and allocation. Changes in metabolite availability altered ATP allocation, biomass accumulation, and transitions between planktonic, aggregative, and biofilm-associated states. Taken together, our results establish a metabolic framework for understanding how environmental conditions regulate the morphology, physiology, and lifestyles of host-associated methanogens, and they may serve as a foundation for the study of other strains.

## Discussion

### Cell morphology and lifestyles in *M. smithii*

Under standard growth conditions in the defined MEP01 medium, *M. smithii* reproducibly exhibited the characteristic morphology and cell division patterns previously described for this species (Miller et al, 1982; Pende et al., 2021; Catlett et al., 2022; Weinberger et al., 2025a; Smith et al., 2026). A distinctive feature of *M. smithii* is an equatorial division septum that constricts the cell into two hemispheres. Septum formation is mediated by a contractile division ring containing FtsZ and SepF, which transiently colocalize at the division plane during binary fission (Pende et al., 2021).

By the time cultures reached late stationary phase (approximately 15 days) and formed cellular aggregates, substrates for methanogenesis had largely been depleted, and the extracellular pH had increased. Although acetate remained detectable in the medium, the increase in extracellular pH to approximately 8.0 would shift the acetate/acetic acid equilibrium toward the deprotonated form, potentially reducing acetate uptake by *M. smithii*, which lacks an identifiable acetate transporter (Samuel et al., 2007; Welte, et al., 2014; Ouboter et al., 2023). We therefore propose that nutrient depletion, together with reduced acetate bioavailability, contributes to the formation of cell aggregates.

Cell aggregates associated with nutrient stress differed morphologically from the biofilms formed under nutrient-rich conditions (**Fig. 1D, F, and H)**, suggesting that distinct mechanisms underlie these alternative lifestyles in *M. smithii*. Cellular aggregates have also been observed during the co-culture of *M. smithii* with the fermentative gut bacterium *Bacteroides thetaiotaomicron* (Catlett et al., 2022).

Furthermore, the resistance of cell aggregates induced by nutrient stress to Triton X-100, DNase I, and proteinase K suggests that extracellular DNA and proteins are unlikely to represent major structural components, in contrast to methanogen biofilms in which these molecules contribute to matrix architecture (Bang et al., 2014; Jasso-Chávez et al., 2015). This observation could implicate uncharacterized proteins that degrade polysaccharides or glycoproteins associated with the S-layer (surface layer) of archaea and other cell surface components such as adhesin-like proteins (Gupta & Seedorf 2024; Tan et al., 2026), or that remodel the cell envelope. An example of this novel proteins is a recently described glycosyl hydrolase that specifically cleaves the peptidoglycan of the archaeal cell wall of *M. smithii* (Smith et al., 2026).

To investigate the molecular basis of the transition to an aggregative lifestyle, we analyzed the gene expression profiles of four independent cultures harvested at the onset of aggregation under standard growth conditions **(Fig. S2)**. The most highly expressed gene was PT320_RS00380, which encodes a diacylglycerol/polyprenol kinase (DAGK/PolK) family member. This enzyme performs two primary functions (Van Horn & Sanders, 2012). Firstly, it provides the foundation for lipid recycling by restoring phosphatidic acid (PA) that has been utilized in the biosynthesis of membrane-derived oligosaccharides (MDOs). Secondly, it facilitates the recycling of diacylglycerol (DAG) phospholipids that are produced during the biosynthesis of lipopolysaccharides (LPS). In bacteria under osmotic stress, there is an increase in MDO production, whereas in plants, polyprenols accumulate in response to hypersaline stress (Baczewska et al., 2014). Furthermore, a polyprenol kinase mutant of *Streptococcus* mutans exhibited a higher sensitivity to acidic conditions, suggesting the involvement of polyprenyl-phosphates in stress response (Lis & Kuramitsu, 2003). In addition, polyprenols have been suggested act as antioxidants that scavenge free radical oxygen species to protect surrounding lipids from peroxidation (Zhang et al., 2015).

Interestingly, a β-class carbonic anhydrase was among the proteins overexpressed. Carbonic anhydrases catalyze the reversible interconversion of CO_2_ and bicarbonate and play important roles in carbon assimilation, acetate uptake and pH homeostasis (Santiago-Martínez et al., 2024). Because formation of cell aggregates occurred as extracellular pH increased and nutrient availability declined, we speculate that increased carbonic anhydrase expression may facilitate bicarbonate-dependent anabolic reactions while maintaining intracellular pH. Alternatively, its induction could reflect a broader range of functions associated with the transition from planktonic growth to an aggregative lifestyle. Further studies are needed to determine its role in adaptation to nutritional stress in *M. smithii*.

We speculate that this cell aggregates phenotype has remained unrecognized because previous studies relied mainly on nutrient-rich media that may mask physiological responses associated with nutrient limitation. Here, the use of a defined medium made it possible to detect these responses and link them directly to changes in nutrient availability.

In contrast, nutrient excess promoted a transition to a biofilm lifestyle (**Fig. 1H**), suggesting that the production of extracellular matrix may function as a carbon sink. Mucosal biofilm formation has previously been reported in *M. smithii* and other methanogens (Bang et al., 2014; Jasso-Chávez et al., 2015), raising the possibility that this lifestyle contributes to host colonization and physical interactions with other members of the gut microbiome. Together with the aggregation phenotypes observed under nutrient limitation, these findings indicate that *M. smithii* exhibits remarkable lifestyle plasticity in response to environmental conditions. The composition of the *M. smithii* biofilm and the regulatory mechanisms governing are the focus of ongoing studies in our laboratory.

### Limited biomass synthesis despite sustained methane production

A notable feature of *M. smithii* physiology was the apparent decoupling of biomass accumulation from methane production. Although final biomass yields remained low, methanogenesis depends mainly on the availability of H_2_, CO_2_ and formate, indicating that energy generation does not necessarily translate into proportional cell proliferation. This observation suggests that a substantial fraction of the energy derived from methanogenesis may be allocated to maintenance functions, stress responses, or lifestyle transitions rather than biomass synthesis. Furthermore, the Msm_MEP model predicts that acetate activation and pyruvate carboxylation are the primary ATP-consuming reactions in *M. smithii*. This finding provides a mechanistic explanation for the strong physiological effects of acetate availability despite its limited consumption, as acetate assimilation imposes a substantial energetic burden on the cell (**Table 2**).

Based on our findings and the genome (Samuel et al., 2007), we propose a revised model of the hydrogen-dependent CO_2_-reducing methanogenesis in *Methanobrevibacter smithii* in which formate and acetate are integral components of the metabolic network, influencing methanogenesis, carbon assimilation, and energy allocation (**Fig. 6**).

Our results also suggest that formate and hydrogen are not functionally interchangeable in *M. smithii*, despite the common assumption that formate can substitute for H_2_ during methanogenesis. While formate oxidation generates reduced cofactor F_420_ (F_420_H_2_) through the activity of formate dehydrogenase (Grinter & Greening, 2021; Holden & Sistu, 2023; Halim et al., 2024), H_2_ primarily contributes reducing power through hydrogenases involved in the generation of reduced ferredoxin (Fdr_ed_), either directly (energy-converting hydrogenase, Eha) or via electron bifurcation (methylviologen-reducing hydrogenase/ heterodisulfide reductase complex, Mvh-HdrABC) (Welte & Deppenmeier 2014; Wagner et al., 2017). Because these electron carriers participate in distinct biochemical reactions, formate and hydrogen support different aspects of methanogen metabolism. In other hydrogenotrophic methanogens, H_2_ and CO_2_ limitation alters the intracellular redox state by changing the ratios of reduced and oxidized cofactors, including F_420_H_2_/F420 and NADH/NAD^+^(Xue et al., 2025) Similar shifts in redox balance may contribute to the physiological and morphological responses observed in *M. smithii* under nutrient-limited conditions.

### Acetate as a biosynthetic precursor and modulator despite its limited consumption

Previous isotope-labeling experiments using 0-10 mM of 2-^13^C-acetate (methyl carbon, CH_3_-) and 1-^13^C-acetate (carboxyl carbon, COO⁻) demonstrated that acetate serves as a precursor for the synthesis of acetyl-CoA and biomass (Choquet et al., 1994a; Choquet et al., 1994b; Moissl-Eichinger et al., 2026) through the activity of an AMP-forming acetyl-CoA synthetase (Acs, *MSM0330* gene) (Samuel et al., 2007). The isotopic label was subsequently detected in ribose, hexoses, purines, pyrimidines and serine, indicating incorporation of acetate-derived carbon into central anabolic pathways.

The additional carbon required for the synthesis of these metabolites is likely supplied by CO_2_ or bicarbonate through reactions involving carbonic anhydrase (Cab *MSM0654, MSM1223*) and pyruvate:ferredoxin oxidoreductase (PFOR, *MSM0557–60*). Subsequent ATP-dependent conversion of pyruvate to phosphoenolpyruvate by PEP synthase (PpsA, *MSM0988*) initiates gluconeogenesis and the production of carbohydrates and other biomass precursors (**Fig. 6**). Consistent with this pathway, the Msm_MEP model identified pyruvate carboxylation and gluconeogenesis as major ATP-consuming processes during biomass synthesis. The functions of these enzymes have been described in other methanogens and appear to be conserved in the *M. smithii* genome (Santiago-Martínez et al., 2016, 2024; Richter et al., 2024; Bonitatibus et al., 2025).

The requirement of *M. smithii* for exogenous acetate (**Fig. 6**) has been attributed to its limited capacity to assimilate CO_2_ through the carbon monoxide dehydrogenase/acetyl-CoA synthase (Cdh/ACS) complex (Lessner et al., 2006; Rother et al., 2006; Biester et al., 2024). Unlike many methanogens and acetogens that grow on CO_2_, formate, or acetate*, M. smithii* lacks key components of this pathway and therefore depends on external acetate as a precursor for acetyl-CoA biosynthesis (Bott et al., 1985; Adam et al., 2018; Nagoya et al., 2021; Garcia et al., 2022).

The inability of *M. smithii* to perform acetoclastic methanogenesis can be partially explained by the absence of the Cdh/ACS complex and key energy-conserving electron transfer systems, including the ferredoxin:NAD⁺ oxidoreductase (Rnf) complex and the F_420_H_2_ dehydrogenase (Fpo) complex. In acetoclastic methanogens, these systems participate in electron transfer from acetate-derived intermediates to membrane-associated energy conservation pathways that generate ion gradients for ATP synthesis (Welte & Deppenmeier, 2014; Downing et al., 2026).

Reported acetate concentrations in the human colon commonly range between 20 and 80 mM, whereas formate concentrations are generally lower and more variable, often in the 0.1–5 mM range (Miller & Wolin, 1986; Hansen et al., 2011; Malat et al., 2024; Moissl-Eichinger et al., 2026). Consequently, the acetate concentrations used in this study are physiologically relevant, while the elevated formate concentrations likely represent the upper range of availability that may occur in localized fermentation hotspots or under conditions associated with gastrointestinal, neuropsychiatric, and neurodevelopmental disorders.

In our experiments, although only small amounts (nmol) of acetate were consumed, the availability of acetate exerted pronounced effects on metabolism, suggesting that acetate may also influence the physiology of *M. smithii* through regulatory mechanisms or as signal. In bacteria, elevated acetate concentrations frequently promote lysine acetylation of metabolic enzymes, with AMP-forming acetyl-CoA synthetase (AMP-Acs) representing one of the best-characterized targets. Acetylation of AMP-Acs inhibits its catalytic activity and modulates carbon flux through central metabolism (Wolfe, 2005).

Both AMP-Acs and the formate dehydrogenase subunit FdhA of *M. smithii* contain several lysine residues corresponding to known acetylation targets in bacteria (Zhang et al., 2020). Furthermore, the *M. smithii* genome encodes both a putative lysine acetyltransferase (*MSM0893*) and a deacetylase (*MSM1087*), indicating that reversible protein acetylation may occur in this methanogen. These observations raise the possibility that acetate availability directly influences enzyme activity and metabolic fluxes through post-translational regulation. Although this hypothesis remains to be experimentally validated, it provides a plausible mechanism by which small changes in acetate availability could exert disproportionately large effects on growth, methanogenesis, and cellular physiology.

Overall, these findings suggest that acetate availability regulates methanogen physiology through a combination of biosynthetic, energetic, and potentially post-translational mechanisms. This helps explain why small changes in acetate concentration produce disproportionately large effects on cellular growth, metabolism, and lifestyles.

### Energetic costs limit biomass production in *M. smithii*

Our Msm_MEP model also predicted that acetate activation, pyruvate carboxylation and gluconeogenesis are among the major ATP-consuming reactions in *M. smithii* (**Table 2**, **Fig. 6**). As extracellular pH increased during late stationary phase, the concentration of protonated acetate likely decreased, potentially limiting acetate uptake despite its continued presence in the medium. This mechanism provides a potential explanation for the strong physiological responses observed under acetate-limited conditions.

Furthermore, changes in pH during growth are expected to alter the partitioning of inorganic carbon among gas-phase CO_2_, dissolved CO_2_, and bicarbonate (HCO_3_⁻) in MEP01 medium. Increased bicarbonate availability could support PFOR-mediated carboxylation reactions (Bonitatibus et al., 2025), thereby contributing additional carbon required for the synthesis of acetyl-CoA-derived metabolites from exogenous acetate. This mechanism is consistent with observations from fermentation systems in which pH regulates inorganic carbon speciation and availability (Flores et al., 2021).

Our Msm_MEP model simulations provided a mechanistic explanation for the distinct physiological responses observed under no formate and low-acetate conditions. In the absence of formate, methane production and ATP generation were substantially reduced. Because Growth-Associated Maintenance (GAM; 50.2 mmol ATP gDW^-1^) scales with growth rate and was included in the ATP allocated to growth; even so, growth accounted for <1% of the ATP produced in the absence of formate (**Table 3**). This response is consistent with methanogenesis-derived ATP supporting the observed growth rates, suggesting that increased nutrient availability enhances methane production, ATP generation, and ultimately biomass accumulation in *M. smithii*.

Under no-formate limitation, model predictions indicated that essentially all ATP production was required to satisfy maintenance demands at the initial biomass concentration, leaving insufficient energy for biomass synthesis. The ATP derived from methanogenesis far exceeded the amount required for the growth observed under all conditions (Poorter et al., 2007; Costa et al., 2013); the limitation in the case of acetate, and the absence of formate, could be due to the availability of precursors. The condition without formate implies that the cell had more ATP at its disposal than was required for the observed growth rate, although this does not necessarily mean that said ATP covered maintenance requirements. This energetic deficit is consistent with the decline in optical density and suggests that cell aggregation (or cell death) occurred because ATP production could no longer sustain growth and maintenance simultaneously. A similar but less severe pattern was observed under low-acetate condition, where most ATP was predicted to be allocated to maintenance rather than biomass production. These findings support the hypothesis that the physiological effects of formate and acetate limitation arise from distinct metabolic constraints that ultimately converge on cellular energy availability, thereby restricting growth while prioritizing maintenance and survival functions.

These findings have important implications for the interpretation of microbiome data. Methane production is frequently used as a proxy for methanogen abundance, particularly in gastrointestinal disorders and intestinal methanogen overgrowth (IMO). However, our results indicate that elevated methane production may reflect increased substrate availability and methanogenic activity rather than increased methanogen biomass. Consequently, methane emissions alone may provide limited information regarding the abundance, metabolic state, or lifestyle of host-associated methanogens.

Our observations further suggest that *M. smithii* prioritizes ATP production through methanogenesis over biomass accumulation when growing on acetate, representing a physiological strategy that may facilitate persistence in the dynamic and nutrient-variable environment of the human gut. Although acetate availability strongly influenced physiology, only limited acetate consumption was required to support growth, indicating that *M. smithii* is more likely to respond to fluctuations in acetate availability than to substantially alter acetate pools within the gut. In contrast, formate availability strongly influenced growth, methanogenesis, extracellular pH, and cellular organization, suggesting that formate metabolism may be a key determinant of *M. smithii* activity *in vivo*.

We cannot exclude the possibility that *M. smithii* supplements its metabolism with additional substrates that impose lower energetic costs on biosynthesis, including lactate, pyruvate, amino acids, methyl-containing compounds or metabolites derived from the metabolism of amines and choline (Feregrino-Mondragón et al., 2023; Richter et al., 2024; Kashyap et al., 2026). Similarly, *M. smithii* may use previously unrecognized energy-conserving mechanisms to increase ATP yield, including supercomplexes, F_420_H_2_-dependent electron transfer systems, nitrogenases, metallochaperones or accessory proteins, or non-canonical methanogenic pathways. (Johnson et al., 2005; Chanderban et al., 2023; Kohtz et al., 2025; Paul et al., 2026; Segundo-Acosta et al., 2026; Dinh et al., 2026). Regardless of the mechanism, our findings identify nutrient availability as a key determinant of methanogen physiology and raise the possibility that *M. smithii* becomes particularly influential in microbial communities and disease states associated with elevated formate production.

## Conclusions

Formate, hydrogen, and acetate emerged as key determinants of the physiology, morphology, and lifestyle transitions of *Methanobrevibacter smithii*. Our findings show metabolite-dependent transitions between planktonic growth, cellular aggregation, and biofilm formation, highlighting an unexpected degree of physiological plasticity in the dominant methanogenic archaeon of the human gut. Because nutrient availability influences these physiological states, important aspects of methanogen function and ecological interactions cannot be inferred from sequencing data alone.

*M. smithii* appears more likely to respond to fluctuations in acetate and formate availability than to substantially alter their concentrations within the gut ecosystem. Moreover, formate, hydrogen, and acetate are not interchangeable substrates but interactive determinants of physiology, contributing distinct metabolic functions and reduced cofactors (e.g., F_420_H_2_ versus Fd_red_). Methane production was partially uncoupled from biomass accumulation, indicating that elevated methane levels may reflect substrate availability and methanogenic activity rather than methanogen overgrowth. These observations suggest that the physiological state of *M. smithii* may be as important as its abundance when interpreting microbiome data and inferring ecological functions or cross-domain interactions.

Collectively, our experimental and modeling results establish a framework linking nutrient availability to methanogen physiology, energy allocation, and cellular organization. This framework provides new insight into how host-associated methanogens respond to environmental fluctuations and may help explain how archaeal physiology shapes cross-domain interactions within the human microbiome.

## Materials and Methods

### Microbial strain and cultivation conditions

The methanogenic archaeon *Methanobrevibacter smithii* PS (DSM 861, ATCC 35061) was used throughout this study, acquired from the Deutsche Sammlung von Mikroorganismen und Zellkulturen (DSMZ). Cultures were maintained and experimentally grown under strict anaerobic conditions at 37°C with continuous shaking at 110 rpm in the dark. The standard cultivation vessel consisted of 118 mL serum bottles containing a defined liquid medium and a H_2_/CO_2_ (80/20, v/v) gas mixture in the headspace at an initial overpressure of ∼0.8 atm. Gas mixtures (Certified Standard Specification) were supplied by Airgas (Radnor, PA, USA). These baseline conditions are referred to as the standard condition.

### Culture medium preparation

A defined, oxygen-free medium with a minimal and defined chemical composition was used for all cultivations. The MEP01 medium contained the following components per liter: 3.0 g Yeast Nitrogen Base (without amino acids), 2.5 g Sodium Bicarbonate, 3.0 g Sodium Formate, 3.0 g Sodium Acetate Trihydrate, 0.5 g L-Cysteine-HCl, and 1.0 mL of a 0.1% (w/v) Resazurin solution.

Yeast Nitrogen Base (DOT #DSY20040) contains: 5.0 g/L ammonium sulfate, (Nitrogen Sources), 1.0 g/L potassium phosphate monobasic, 0.5 g/L Magnesium sulfate, 0.1 g/L sodium chloride, 0.1 g/L calcium chloride, vitamins (2.0 μg/L biotin, 400 μg/L calcium pantothenate, 2.0 μg/L folic acid, 400 μg/L nicotinic acid, 200 μg/L p-aminobenzoic acid, 400 μg/L pyridoxine HCl, 200 μg/L riboflavin, 400 μg/L thiamine HCl, and 2 mg/L inositol) and trace elements (500 μg/L boric acid, 40 μg/L copper sulfate, 100 μg/L potassium iodide, 200 μg/L ferric chloride, 400 μg/L manganese sulfate, 200 μg/L sodium molybdate, and 400 μg/L zinc sulfate). This base media does not contain amino acids or carbon sources. We performed tests using enzymatic assays and observed that formate, acetate, phosphoenolpyruvate, pyruvate, lactate, glutamate, citrate and glucose were not present in the cell-free medium (Santiago-Martínez et al., 2016; Hernández-Esquivel, Del Mazo-Monsalvo et al., 2025).

The medium was prepared according to the following procedure to ensure anaerobiosis: All components, except L-Cysteine-HCl, were weighed and dissolved in the final volume of deionized water. The solution was transferred into an anaerobic chamber and was stirred on a magnetic stirrer with an aquarium pump for gas circulation for approximately 16 hours to remove oxygen and CO_2_. The pH was adjusted to approximately 7.2 using drops of a fresh 1:10 dilution of HCl until all precipitates dissolved. L-Cysteine-HCl (0.5 g/L) was added as a reducing agent, and the solution was stirred for 1 hour until it turned from purple to hot pink, indicating a reduced state (absence of oxygen).

The medium was transferred into serum bottles at storage volume (80 mL). Bottles were sealed with rubber stoppers and aluminum crimp caps. The headspace was flushed with the H_2_/CO_2_ gas mixture, and the bottles were autoclaved at 120°C for 30 minutes. After cooling to room temperature, the headspace was flushed again with the H₂/CO₂ mixture. Immediately before inoculation, 0.025% (w/v) Na_2_S was added, and the headspace was flushed once more to establish the target initial overpressure.

Serum bottles of different capacities (118 mL for experiments; 1 L and 2 L for culture expansion) were prepared using a standardized two-step sterilization and gassing procedure. Empty bottles were sealed with butyl rubber stoppers and aluminum crimp caps, flushed with the H_2_/CO_2_ mixture (80/20, v/v), and autoclaved at 120°C for 30 minutes. After cooling to room temperature, the headspace was flushed again with the same gas mixture to ensure anaerobiosis and establish the desired overpressure. These generated sterile, pre-gassed bottles ready to receive liquid volumes as needed.

### Experimental setup and inoculation

For each experimental batch, a single 1 L master culture (OD_600_ ≥ 0.1, late exponential phase) was used to inoculate all experimental bottles (See supplementary material). Using anaerobic syringe techniques, the appropriate volume of master culture was transferred into each prepared serum bottle to achieve a standardized initial optical density of ∼0.040 at 600 nm. This target range was selected based on preliminary optimization trials as it provided the best balance between a well-defined lag phase, consistent growth kinetics, and maximum dynamic range to detect treatment effects. Typical inoculation volumes were approximately 6 mL per 100 mL experimental bottle, with proportional adjustments for larger vessels or bottles.

A consistent two-step gassing procedure was employed at inoculation: 1) Immediately before inoculation, the headspace was flushed with the H_2_/CO_2_ mixture. 2) Immediately after inoculation, the headspace was flushed again to establish a defined initial overpressure of approximately 0.8 atm (≈ 11.8 PSI), verified using a digital manometer.

### Standard and variant conditions

The primary experimental condition, designated as *Standard*, consisted of cultures grown in the defined medium under H_2_/CO_2_ (80/20) atmosphere at 0.8 atm overpressure.

To investigate specific metabolic dependencies, the following variant conditions were established, each modifying a single parameter from the Standard condition. When a sodium salt was omitted, an equimolar amount of NaCl was added to maintain sodium ion concentration: A) <u>No-formate:</u> Sodium formate omitted from the medium, B) <u>Argon</u>: H_2_/CO_2_ replaced by Ar/CO_2_ (80%/20%), C) <u>Tungsten</u>: Sodium tungstate added at 0.05, 0.5, 5, 50, and 500 µM, D) <u>Low-acetate:</u> Sodium acetate omitted from the medium.

All conditions were tested in at least triplicate within each experimental batch. To ensure reproducibility, each condition was evaluated across at least three independent batches (derived from different master cultures), with 3–5 replicate bottles per condition per batch. Concurrent controls (3–5 bottles of standard condition) were included in every batch.

### Experimental culture configuration

Experimental cultures were conducted in 100 mL serum bottles with a 20 mL working volume (A/V ≈ 1 cm^2^/mL), as this configuration represents the standard condition. Cultures in 1 L and 2 L bottles (100 mL and 150 mL working volumes, respectively) were used when larger sample volumes were required for specialized analyses.

### Sampling for analytical monitoring

We use gas chromatography analysis to quantify gases (methane, hydrogen, and carbon dioxide) as direct parameters for assessing active microbial metabolism. To minimize disturbances to the anaerobic cultures, a strict order was followed for all measurements at each sampling time point: headspace gas samples were collected first (200 µL), followed by headspace pressure measurement (which requires a brief gas flow through the digital manometer), and finally liquid samples (1 mL) for determination of optical density at 600 nm (OD_600_). This sequence preserves the headspace composition for gas chromatography while ensuring that pressure readings reflect the undisturbed culture status and places the most invasive step (liquid removal) last. Sampling frequency was empirically optimized during the study. Initial experiments with daily sampling (serial measurements on the same bottles) revealed that frequent manipulation altered culture performance. Therefore, for routine screening, bottles were sampled with low frequency (≤6 time points per bottle). For detailed kinetic characterization, higher sampling frequencies (>6 time points) were employed, but using independent biological replicates for each time point to avoid cumulative disturbance. All time points were randomized across treatments to prevent systematic bias.

### Growth monitoring

Growth was tracked by measuring optical density at 600 nm (OD_600_) using a UV-1600PC spectrophotometer. A single 1 mL liquid aliquot was aseptically removed from each serum bottle. This aliquot was first used for OD measurement, then centrifuged. The resulting supernatant was used for pH measurement and subsequent enzymatic quantification of acetate and formate concentrations.

### Headspace pressure and gas composition

Total pressure within serum bottles was monitored using a digital manometer. For gas composition analysis, a 200 µL sample was withdrawn from the headspace. Concentrations of H_2_, CO_2_, and CH_4_ were determined using an Agilent 8890 Gas Chromatography (GC) System. Pressure data were referenced to abiotic controls to establish accurate initial values, with all subsequent changes attributable to biological activity. System integrity tests established an operational lower limit of −0.459 ± 0.048 atm (mean ± SD, n=8 independent determinations), beyond which the risk of back-diffusion or seal compromise increases significantly, providing a natural endpoint for gas-consumption experiments.

### Calculation of gas consumption/production

Partial pressure data were plotted over time and fitted with linear or exponential models as appropriate. The slope of these fits, representing the rate of pressure change, was used to calculate the daily volume of gas consumed or produced. This volume was then converted to total micromoles of each gas using the ideal gas law, allowing quantification of metabolic activity.

### Content of metabolites and coupled enzymatic essays

The content of metabolites (formate, acetate, phosphoenolpyruvate, pyruvate, lactate, glutamate, citrate and glucose) in liquid media was determined by monitoring the reduction/oxidation of NAD(H) or NADP(H) at 340 nm, as previously described (Santiago-Martínez et al., 2016; Hernández-Esquivel, Del Mazo-Monsalvo et al., 2025). Coupled enzymatic assays were performed at 37 °C in a multi-cell spectrophotometer (Agilent, Santa Clara, CA, USA), using commercial enzymes and samples collected at various time points during growth or from cell-free media. For the quantification of formate, commercial formate dehydrogenase from *Candida boidinii* (Sigma-Aldrich #F8649) was used.

### Kinetic modeling of cell growth and metabolism

Cell growth was modeled with a modified Gompertz equation incorporating death (Zwietering et al., 1990). Methane production followed standard Gompertz kinetics. Substrate (H_2_, CO_2_) consumption was described by linear or exponential decay models as appropriate. All fittings were performed in OriginPro 2026, using the equation in **Table 4**, with model selection based on R^2^ and parameters (See supplementary material).

**Table 4.** Equations and variables used for kinetic modeling of cell growth and gas metabolism.

| Equation number | Name | Short name | Equations |
|---|---|---|---|
| Equation 1 | Modified Gompertz (with cell aggregation component) | MG | $y = y_0 + A \cdot \exp\big(-\exp(-r \cdot (x - t_i))\big) \cdot \exp(-\delta \cdot x)$ |
| Equation 2 | Standard Gompertz | SG | $y = A \cdot \exp\big(-\exp(-r \cdot (x - t_i))\big)$ |
| Equation 3 | Maximum specific growth rate | µmax | $\mu_{max} = \frac{A \cdot r}{e}$ |
| Equation 4 | Doubling time | t<sub>d</sub> | $t_d = \frac{\ln(2)}{\mu_{max}}$ |
| Equation 5 | Lag phase duration | λ | $\lambda = t_i - \frac{1}{r}$ |
| Equation 6 | Methane production (Gompertz) |  | $CH_4 = A_{CH_4} \cdot \exp\Big(-\exp\big(-r_{CH_4} \cdot (x - t_{i,CH_4})\big)\Big)$ |
| Equation 7 | Maximum methanogenic rate | µCH<sub>4</sub> | $\mu_{CH_4} = \frac{A_{CH_4} \cdot r_{CH_4}}{e}$ |
| Equation 8 | H<sub>2</sub> consumption (first-order exponential decay) |  | $[H_2] = [H_2]_0 \cdot \exp\big(-k_{H_2} \cdot t\big)$ |
| Equation 9 | H<sub>2</sub> half-life | t<sub>1</sub>/<sub>2</sub> | $t_{1/2} = \frac{\ln(2)}{k_{H_2}}$ |
| Equation 10 | H<sub>2</sub> consumption rate constant relationship | kH<sub>2</sub> | $k_{H_2} = \frac{1}{\tau}$ |
| Equation 11 | CO<sub>2</sub> consumption (linear model) |  | $[CO_2] = m \cdot t + b$ |

### Statistical Analysis

For each experimental batch, treatment effects were evaluated against their concurrently paired controls (Cn) from the same master culture inoculum. This paired approach accounts for day-to-day variability in inoculum activity and handling.

All conditions were tested across at least three independent batches (biological replicates), each derived from a distinct master culture. Within each batch, conditions were tested in at least triplicate (technical replicates). Data are presented as mean ± standard deviation unless otherwise indicated.

### Scanning electron microscopy (SEM)

Samples were collected from each mentioned condition, with 5mL of sample being deposited into 24-well culture plates with circular slides coated in poly-L-lysine. The samples were left to adhere on the slides for approximately 10 minutes, and after removing the supernatant, were rinsed with 2.5 mL of 0.12 M phosphate buffer + 3 mM MgCl_2_, pH 7.4. After this, 2.5 mL of the fixative solution (2% glutaraldehyde + 1.5% paraformaldehyde in 0.12 M phosphate buffer + 3 mM MgCl_2_, pH 7.4) was added to the samples and incubated at 4°C overnight. The fixative was then rinsed off with 2.5 mL 0.12 M phosphate buffer + 3 mM MgCl_2_, pH 7.4, and put on a rotator slow setting for 2 sessions of 30 minutes with the supernatant being removed each time. After the final buffer wash, the samples were rinsed with distilled water and put on the rotator for 30 minutes, slow settings 2 times with supernatant being removed. This was then followed by serial dehydration in ethanol (30%, 50%, 70%, 95%, 100%, 100%) for 30 minutes each, slow settings on the rotator and kept in 100% ethanol overnight. Samples were then rinsed in 100% fresh ethanol, 30 minutes slow setting on rotator the following day prior to critical point drying. The samples were then critical point dried (931GL, Tousimis) and mounted onto SEM stubs using double sided carbon tape. They were sputter coated with 10nm of platinum (CCU-010, Safematic) and examined using a scanning electron microscope (Nova NanoSEM 450, FEI) at 2kV accelerating voltage.

### Curation of Genome-scale metabolic model (GEM)

Our Msm_MEP model was refined from the AGORA2 GEM of *M. smithii* ATCC 35061 reconstruction (Heinken et al., 2023). All changes are documented in supplementary **Table S1**. Gene assignments were mapped to RefSeq locus tags and Uniprot (proteome UP000001992) (Samuel et al., 2007). Reactions lacking gene support were searched by sequence (BLAST) against characterized archaeal enzymes. Reactions that remained unresolved were kept or added if required for biomass production in the defined medium and due to archaeal presence or published methanogen models (Shoaie et al., 2013; Richards et al., 2016; Casini et al., 2023). Reactions were checked for elemental and charge balance against Rhea and ChEBI, and the model was tested for consistency using MEMOTE 0.17.0 (Lieven et al., 2020). The MEMOTE score of Msm_MEP is 87.2%.

### Constraint-based modeling

Simulations used Msm_MEP in COBRApy 0.31.1 (Ebrahim et al., 2013) with GNU Linear Programming Kit (GLPK) package (http://www.gnu.org/software/glpk/glpk.html). Uptake was limited to medium components (Thiele & Palsson, 2010; Orth et al., 2010), with formate or H_2_ uptake closed for the no-formate and no-H_2_ conditions and the low-acetate condition used the standard medium bounds, excluding acetate Na^+^ translocation by Mtr (1.7 Na⁺ per CH_4_) and the A_1_A_o_ ATP synthase (4 Na+ per ATP) gives 0.425 ATP per CH_4_ (Lienard et al., 1996; Poorter et al., 2007; Richards et al., 2016; Costa et al., 2013). Growth-associated maintenance (GAM) was set to 50.2 mmol ATP gDW^-1^ (Shoaie et al., 2013; Goyal et al., 2015). Specific growth and CH_4_ production rates were calculated from Gompertz model fits at selected time points. For the standard condition, rates were estimated at the maximum CH_4_ production rate (day 3.90), the OD_600_ inflection point (day 4.61), and day 10. For the no-formate condition, rates were estimated at the maximum CH_4_ production rate (day 6.94). For the low-acetate condition, maximum OD_600_ and the linear CH_4_ production rate were used. For the no-H_2_ condition, the initial OD_600_ was used, with the growth rate constrained to zero. Biomass dry weight was estimated from the protein/OD_600_ calibration (**Fig. 2C**), assuming 0.564 g protein per g dry weight (gDW), based on the model biomass composition. Each selected time point was treated as a quasi-steady state (Varma & Palsson 1994; Mahadevan et al. 2002). At each timepoint, growth and CH_4_ fluxes were fixed to the measured values and ATP maintenance flux was maximized with GAM removed (Pirt 1965; Richards et al., 2016; Lachance et al., 2019). ATP not accounted for growth (NGAM) was calculated as this ATP surplus minus GAM x µ (i.e. ATP not accounted for by modelled growth. ATP costs of biomass synthesis were obtained by Parsimonious Flux Balance Analysis (pFBA) at maximal growth with NGAM = 0, With CH_4_ fixed at Standard peak rate (29.53mmol gDW/h) (Lewis et al., 2010). ATP used by adenylate kinase was assigned to AMP-forming reactions (2 ATP equivalents per AMP), and Na⁺ re-entering the cell other than through the ATP synthase was counted as “ATP is not produced” (4 Na⁺ per ATP). The model, medium, and code will be made available as described in Data Availability.

## Acknowledgments

This work was supported by start-up funds and an Urgent Seed Grant (USG26) awarded to MGS-M through the University of Connecticut (UConn), with support from the Department of Molecular and Cell Biology, the College of Liberal Arts and Sciences, and the Office of the Vice President for Research (OVPR).

We thank Dr. Eran Agmon (UConn Health) for his valuable guidance in the development of the genome-scale metabolic modeling tools presented in this work. We also thank Dr. Xuanhao Sun and Dr. Maritza Abril, from the UConn Bioscience Electron Microscopy Laboratory, for their support with microscopy.

We are grateful to the scientists with disabilities who contributed to this work and to the DisabledInSTEM program for its mentorship, community support, and commitment to advancing accessibility and inclusion in STEM.

We thank the founders, organizers, and members of the Archaea Power Hour and Archaea.Bio initiatives for creating a vibrant global community that highlights archaeal research and fosters the open exchange of protocols, tools, and ideas. This article is lovingly dedicated to all the scientists whose curiosity, dedication, and discoveries have advanced the study of archaea over the years.

## Author Contributions

**Isis Del Mazo-Monsalvo:** Conceptualization, Data curation, Formal analysis, Investigation, Methodology, Validation, Visualization, Writing – original draft, Writing – review and editing.

**Alexander N. Poulter:** Conceptualization, Data curation, Formal analysis, Investigation, Methodology, Software, Validation, Visualization, Writing – review and editing.

**Cesar Ivan Ovando-Ovando:** Data curation, Formal analysis, Investigation, Methodology, Software, Validation, Visualization, Writing – review and editing.

**Michel Geovanni Santiago-Martínez:** Conceptualization, Resources, Data curation, Formal analysis, Investigation, Methodology, Validation, Visualization, Writing – original draft, Writing – review and editing.

## Data availability

All data necessary to evaluate the conclusions of this study are included in the main text and/or the Supplementary Materials. The Msm_MEP genome-scale metabolic model (GEM), associated code, and supporting files will be deposited in a publicly accessible GitHub repository once this manuscript has been published following a peer-review process.

## Statement on the use of artificial intelligence (AI) tools

During development of the GEMs, Claude, Copilot, ChatGPT and other large language model–based tools were used to assist with coding and data-querying tasks. All AI-generated code and associated outputs were manually reviewed and validated using the methods described in the cited references.

## Supplementary Material

Additional results are shown in Table S1, Table S2 Figure S1 and Figure S2. Detailed methods and graphical representation of cell culture protocol in Figure S3 and Figure S4. Table S3 contains details of the Msm_MEP GEM model. These files will be deposited in a publicly accessible GitHub repository once this manuscript has been published following a peer-review process.

